# EMT changes actin cortex rheology in a cell-cycle dependent manner

**DOI:** 10.1101/2020.12.15.422849

**Authors:** K. Hosseini, A. Frenzel, E. Fischer-Friedrich

## Abstract

The actin cortex is a key structure for cellular mechanics and cellular migration. Accordingly, cancer cells were shown to change their actin cytoskeleton and their mechanical properties in correlation with different degrees of malignancy and metastatic potential. Epithelial-Mesenchymal transition (EMT) is a cellular transformation associated with cancer progression and malignancy. To date, a detailed study of the effects of EMT on the frequency-dependent viscoelastic mechanics of the actin cortex is still lacking. In this work, we have used an established AFM-based method of cell confinement to quantify the rheology of the actin cortex of human breast, lung and prostate epithelial cells before and after EMT in a frequency range of 0.02 – 2 Hz. Interestingly, we find for all cell lines opposite EMT-induced changes in interphase and mitosis; while the actin cortex softens upon EMT in interphase, the cortex stiffens in mitosis. Our rheological data can be accounted for by a rheological model with a characteristic time scale of slowest relaxation. In conclusion, our study discloses a consistent rheological trend induced by EMT in human cells of diverse tissue origin reflecting major structural changes of the actin cytoskeleton upon EMT.

**Significance statement:** The actin cortex is a key structure for cellular mechanics and cellular migration. Correspondingly, migratory cancer cells were shown to change their mechanical properties to a softer phenotype. EMT is a cellular transformation associated with cancer progression and malignancy. To date, a detailed study of the effects of EMT on the mechanics of the actin cortex is still lacking. In this work, we provide such a study for human breast, lung and prostate epithelial cells in dependence of the cell cycle stage. We observe a softening of the actin cortex in interphase but stiffening in mitosis upon EMT. In conclusion, our study discloses a consistent mechanical trend induced by EMT in human cells of diverse tissue origin.

## Introduction

Many cells deform to function. Consequently, it is not surprising that cell mechanics is becoming more recognised to be a major factor in multiple diseases, such as cancer and cardiovascular, liver, and renal diseases [1, 2]. The actin cortex is a key contributor to passive material stiffness of cells, cellular ability to generate active forces and cell shape regulation [1, 3]. Hence, understanding actin cortex rheology is vital in understanding cell shape regulation.

The actin cortex is a thin layer of a densely cross-linked actin polymer network beneath the plasma membrane of animal cells. It is a complex material with time-dependent viscoelastic mechanics [1, 3]. Further, it is subject to an intrinsically generated active contractile tension; this tension stems from the action of molecular motor proteins (myosins), which act as active force dipoles in the actin polymer meshwork [3]. Molecular constituents of the actin cortex are subject to dynamic turnover (e.g. polymerisation and depolymerisation) - a feature that inevitably contributes to elastic relaxation processes in the material as well as to a time-limited shape memory [3].

Epithelial cells line the surface of organs and internal cavities in animals [4]. If epithelial cells have undergone carcinogenic transformations, they constitute a type of cancer called carcinoma. Carcinoma make up the majority (~ 85%) of malignant tumors in humans [5]. The Epithelial-Mesenchymal transition (EMT) is a cellular transition that converts polarised stationary epithelial cells into migratory mesenchymal cells. It is known to be a hallmark of cancer progression and malignancy in carcinoma [6–8]. EMT is commonly linked to early steps in metastasis promoting cell migration and cancer cell invasiveness [6–9]. Furthermore, EMT was shown to give rise to major changes in the molecular regulation of the actin cortex; the cytoskeletal regulator proteins RhoA and Rac1 undergo characteristic activity changes upon EMT indicating EMT-induced changes in the polymerisation process and the structure of the actin cytoskeleton [10–13].

While mechanical changes of cancer cells have been documented in many studies [10, 14–20], a detailed study of EMT-induced rheological changes of the actin cortex is still lacking. Here, we present such a rheological characterisation in dependence of malignant potential, EMT and cell-cycle stage. We utilise an established cell mechanical assay based on cell confinement with a wedged cantilever of an atomic force microscope (AFM) [10, 21, 22]. In this measurement setup, initially round mitotic or suspended interphase cells are uniaxially compressed between two parallel surfaces and exposed to time-periodic cell height oscillations [10, 22] (Fig. 1a,b). As was shown before, the resulting mechanical response of the cell in this setup is dominated by the actin cortex, see Fig. 4 in [22] and Fig. S7 in [10]. Correspondingly, we use the measured AFM force readout to estimate active cortical tension and the complex elastic modulus of the actin cortex in dependence of frequency in interphase and mitosis in epithelial and EMT-transformed mesenchymal cells [10, 21, 22].

**Figure 1.**
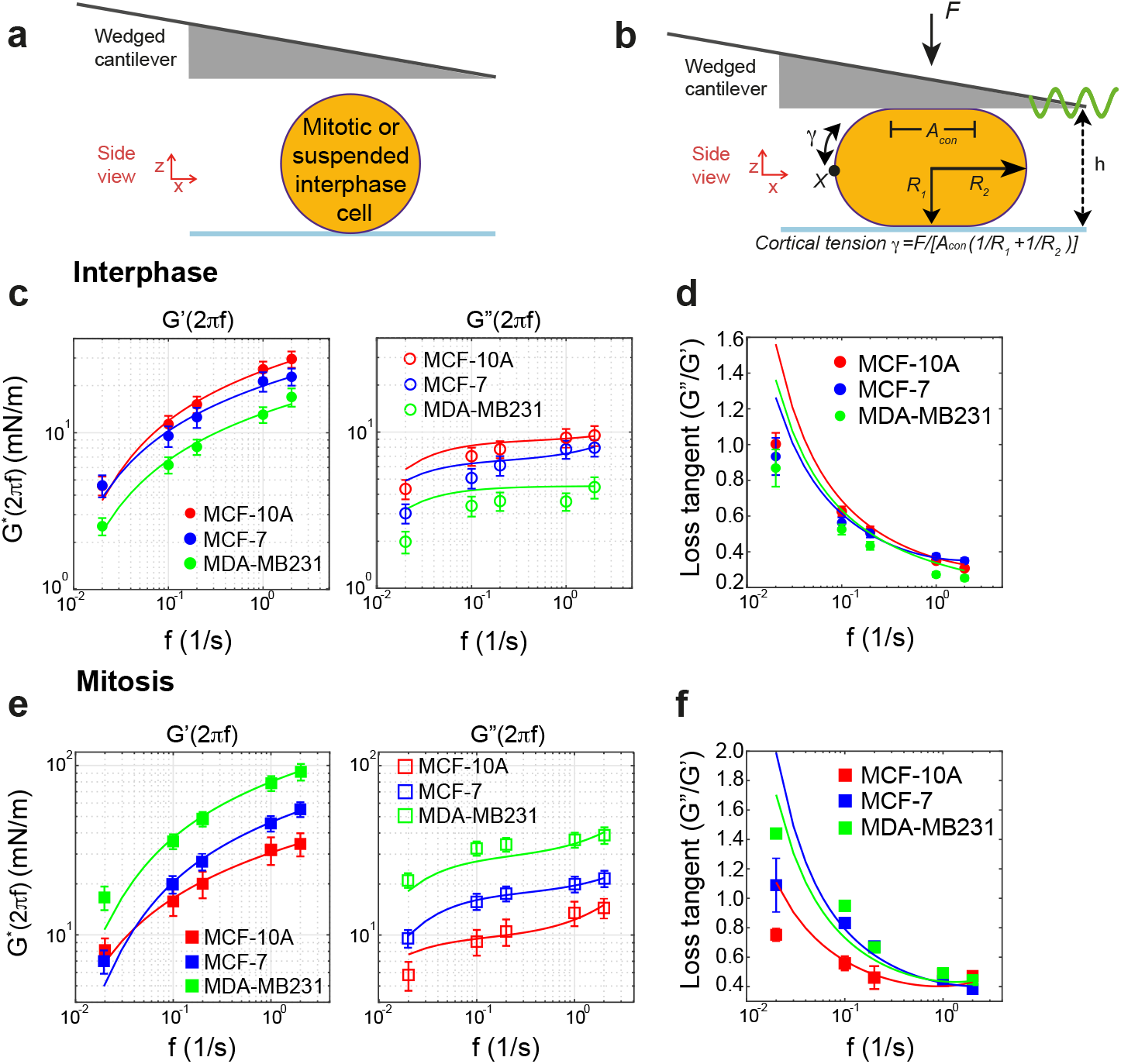
Mechanical phenotyping of the actin cortex for two breast-epithelial cell lines with epithelial phenotype: MCF-10A and MCF-7, and one with mesenchymal-like phenotype: MDA-MB231. Measurements were performed on interphase cells in suspension and STC-arrested mitotic cells. a-b) Schematic of experimental setup. Non-adherent interphase or mitotic cells (yellow sphere) are confined by a wedged AFM cantilever. Dynamic cantilever height changes deform the cell around a mean confined reference shape while recording cellular force response through AFM cantilever deflection. c, e) Averaged twodimensional complex elastic moduli of suspended interphase cells (c) and mitotic cells (e) of cell lines MCF-10A, MCF-7 and MDA-MB231 (Storage modulus: filled symbols, loss modulus: open symbols). Solid lines represent fits of our rheological model (see Eqn. (2) and (3)). d,f) Loss tangents of interphase cells (d) and mitotic cells (f) corresponding to data in panel c and panel e. Solid lines represent loss tangents as predicted by our rheological model with parameters obtained from fitting of storage and loss moduli as depicted in panels c and e. Respective fit parameters were: MCF-10A interphase: *K_h_* = 5.629 mN/m, *τ_max_* = 9.139 s and *η* = 0.3 Pas, MCF-7 interphase: *K_h_* = 4.19 mN/m, *τ_max_* = 8.37 s and *η* = 0.4 Pas, MDA-MB231 interphase: *K_h_* = 2.87 mN/m, *τ_max_* = 10.16 s and *η* = 0.01 Pas, MCF-10A mitosis: *K_h_* = 6.129 mN/m, *τ_max_* = 23.242 s and *η* = 0.431 Pas, MCF-7 mitosis: *K_h_* = 11.225 mN/m, *τ_max_* = 9.445 s and *η* = 0.3232 Pas and MDA-MB231 mitosis: *K_h_* = 18.6543 mN/m, *τ_max_* = 11.6887 s and *η* = 0.9314 Pas. Number of cells analysed: interphase: MCF-10A n=32, MCF-7 n=39, MDA-MB231 n=24, mitosis: MCF-10A n=32, MCF-7 n=38 and MDA-MB231 n=41. Error bars indicate the standard error of the mean. Normalised confinement heights chosen during respective measurements are given in Table I.

**Figure 2.**
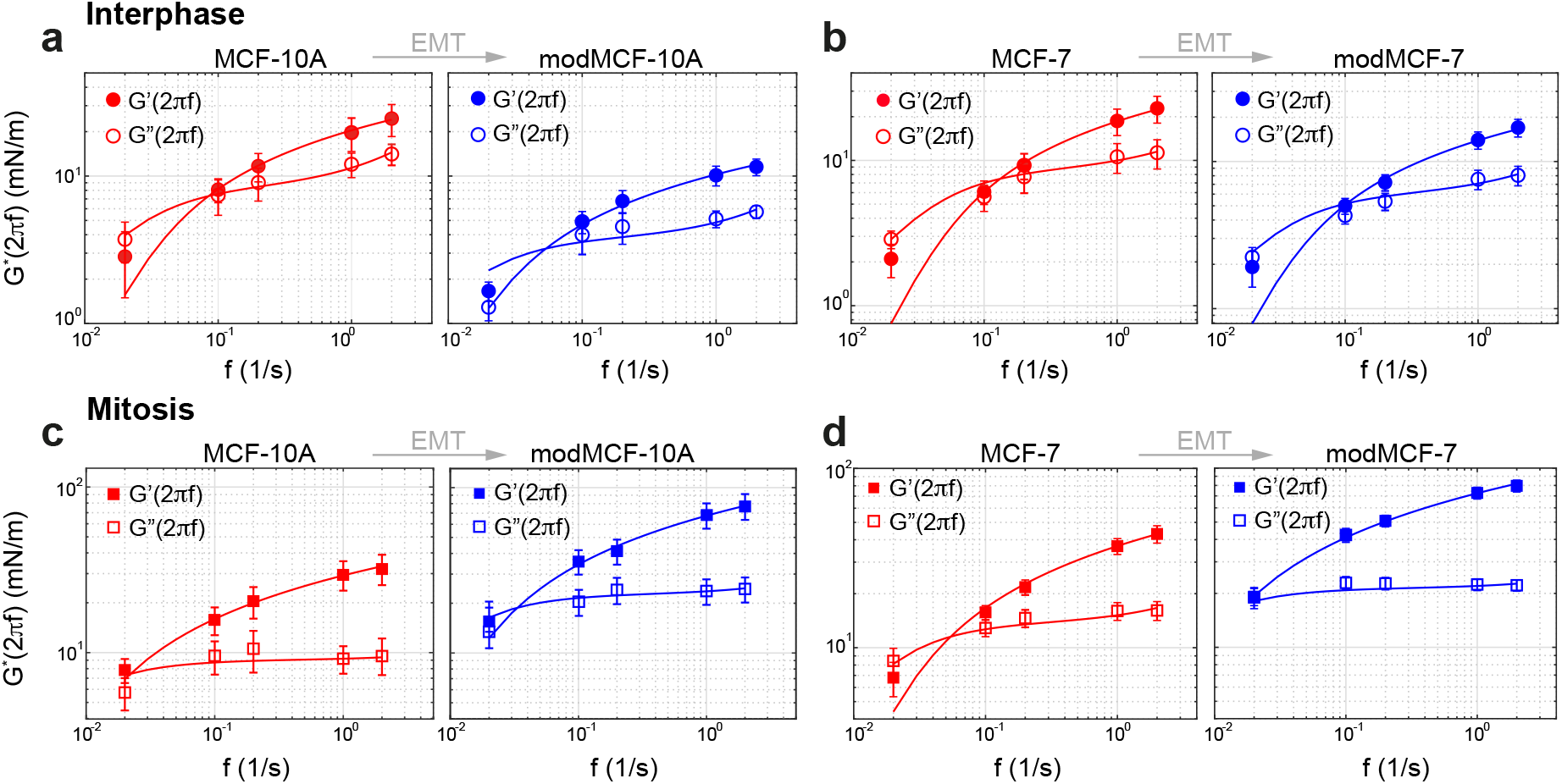
EMT-induced actin cortex mechanical changes of MCF-10A and MCF-7 cells. a-b) Averaged two-dimensional complex elastic moduli of suspended interphase MCF-10A and MCF-7 cells upon EMT. c-d) Averaged two-dimensional complex elastic moduli of MCF-10A and MCF-7 cells in mitotic arrest before and after EMT. (Post-EMT cells are referred to as modMCF-10A and modMCF-7, respectively.) In a-d, solid lines represent fits of our rheological model (Eqn. (2) and (??): MCF-10A interphase: *K_h_* = 5.38 mN/m, *τ_max_* = 7.067 s and *η* = 0.484 Pas, modMCF-10A interphase: *K_h_* = 2.4318 mN/m, *T_max_* = 10.775 s and *η* = 0.17 Pas, MCF-7 interphase: *K_h_* = 3.85 mN/m, *τ_max_* = 4.431 s and *η* = 0.226 Pas, modMCF-7 interphase: *K_h_* = 3.9 mN/m, *τ_max_* = 5.63 s and *η* = 0.171 Pas, MCF-10A mitosis: *K_h_* = 5.77 mN/m, *τ_max_* = 25.27 s and *η* = 0.25 Pas, modMCF-10A mitosis: *K_h_* = 16.12 mN/m, *τ_max_* = 19.83 s and *η* = 0.30 Pas, MCF-7 mitosis: *K_h_* = 8.827 mN/m, *τ_max_* = 10.39 s and *η* = 0.23 Pas and modMCF-7 mitosis: *K_h_* = 13.66 mN/m, *τ_max_* = 31.9 s and *η* = 0.1 Pas. Number of cells analysed: interphase: MCF-10A n=16, modMCF-10A n=18, MCF-7 n=20, modMCF-7 n=18, mitosis: MCF-10A n=12, modMCF-10A n=13, MCF-7 n=23 and modMCF-7 n=22. Error bars indicate standard error of the mean. Normalised confinement heights chosen during respective measurements are given in Table II.

**Figure 3.**
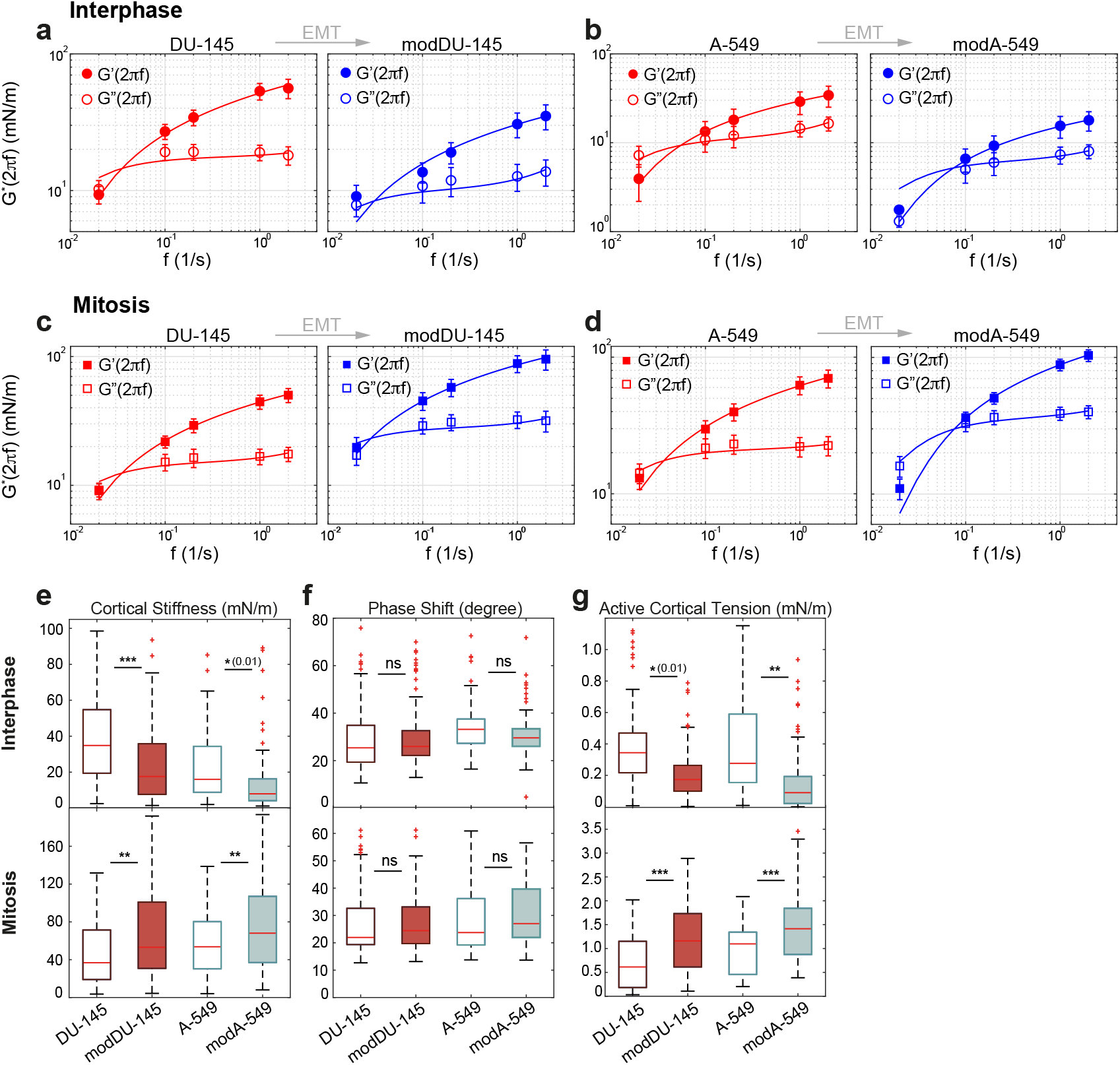
EMT-induced actin cortex mechanical changes of DU-145 and A549 cells. a-b) Averaged two-dimensional complex elastic moduli of suspended interphase DU145 and A-549 cells upon EMT. c-d) Averaged two-dimensional complex elastic moduli of DU145 and A-549 cells in mitotic arrest before and after EMT. In panels a-d, solid lines represent fits of our rheological model (Eqn. (2) and (3)) to measured storage and loss moduli. e-g) Boxplots of cortical stiffness |*G*\*| (e), phase shift (f) and active cortical tension (g) measured for suspended interphase cells (top row) and cells in mitotic arrest (bottom row) before and after EMT at frequency *f* = 1 Hz. (Post-EMT cells are referred to as modDU-145 and mod-A-549, respectively. Fitting measured storage and loss moduli in panels a-d to our rheological model, we obtained the following fit parameters: DU145 interphase: K_h_ = 11.2 mN/m, *τ_max_* = 15.91 s and *η* = 0.102 Pas, modDU145 interphase: *K_h_* = 6.43 mN/m, *τ_max_* = 18.18 s and *η* = 0.331 Pas, A-549 interphase: *K_h_* = 7.05 mN/m, *τ_max_* = 10.44 s and *η* = 0.467 Pas, modA-549 interphase: *K_h_* = 3.95 mN/m, τ_mαx_ = 7.61 s and *η* = 0.156 Pas, DU145 mitosis: K_h_ = 9.6 mN/m, *τ_max_* = 16.11 s and *η* = 0.227 Pas, modDU145 mitosis: *K_h_* = 17.85 mN/m, *τ_max_* = 19.85 s and *η* = 0.409 Pas, A-549 mitosis: *K_h_* = 13.51 mN/m, *τ_max_* = 15.47 s and *η* = 0.147 Pas and modA-549 mitosis: *K_h_* = 22.96 mN/m, *τ_max_* = 7.44 s and *η* = 0.383 Pas. Number of cells analysed (a-d): interphase: DU145 n=24, modDU145 n=22, A-549 n=22, modA-549 n=24, mitosis: DU145 n=18, modDU145 n=17, A-549 n=24 and modA-549 n=24. Number of cells measured at f = 1 Hz (e-g): interphase: DU145 n=36, modDU145 n=36, A-549 n=35, modA-549 n=39, mitosis: DU145 n=34, modDU145 n=34, A-549 n=34 and modA-549 n=29. Error bars indicate standard error of the mean. n.s.: *p* > 0.05, *: *p* < 0.05, **: *p* < 0.01, * * *: *p* < 0.001). Normalised confinement heights chosen during presented measurements are given in Table II.

**Figure 4.**
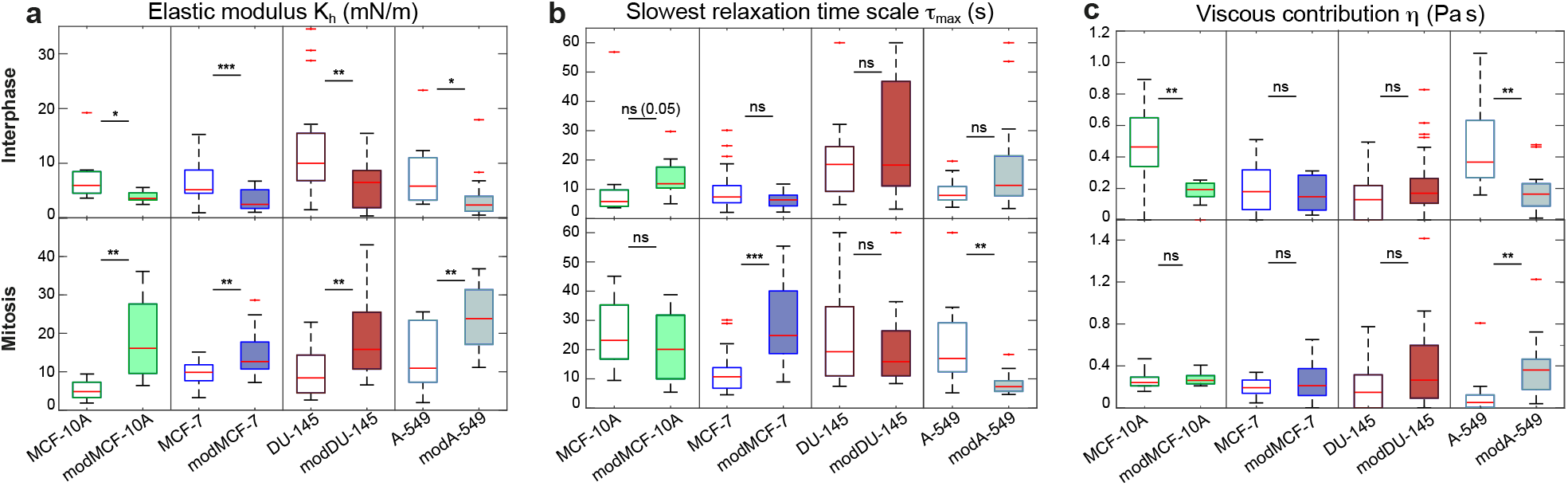
a-c) Boxplots indicating the distribution of rheological fitting parameters (a) *K_h_*, (b) *τ_max_* and (c) *η* for measured MCF-10A, MCF-7,DU-145, A-549 interphase and mitotic cells before and after EMT. (Post-EMT cells are referred to as modMCF-10A, modMCF-7, modDU-145 and mod-A-549, respectively. Number of cells analysed (a-d): interphase: MCF-10A n=16, modMCF-10A n=18, MCF-7 n=20, modMCF-7 n=18, DU-145 n=24, modDU-145 n=22, A-549 n=22, modA-549 n=24, mitosis: MCF-10A n=21, modMCF-10A n=23, MCF-7 n=23, modMCF-7 n=22, DU-145 n=18, modDU-145 n=17, A-549 n=24 and modA-549 n=24. n.s.: *p* > 0.05, *: *p* < 0.05, **: *p* < 0.01, * * *: *p* < 0.001.)

## Results

In the first set of experiments, we investigated three breast epithelial cell lines with increasing degree of malignant potential. The first cell line, MCF-10A, represents healthy breast epithelial cells. The second cell line, MCF-7, are breast carcinoma cells with epithelial phenotype. The third cell line, MDA-MB231, are also breast carcinoma cells that have undergone EMT and exhibit consequently a mesenchymal-like phenotype [14, 23–25]. These three cell lines exhibit a graded increase of migration ability and invasiveness, with MCF-10A showing the smallest, MCF-7 an intermediate and MDA-MB231 the largest migration ability [26–28]. Correspondingly, a graded increase of malignant potential is associated to these cell lines [14, 15, 23]. Here, we performed a frequency-dependent analysis of the rheology of the actin cortex of all three cell lines, to see whether altered malignant potential is reflected in changes of cortical mechanics.

Results of our rheological analysis of the three cell lines are presented in Fig. 1c-f. We find that the frequency dependence of elastic moduli is qualitatively similar to that of the cortical elastic moduli observed for the actin cortex of HeLa cells [22]; at high frequencies, the storage modulus *G*′ clearly dominates (Fig. 1c,e). However, in the low frequency regime, storage and loss modulus are of similar magnitude, indicating a higher fluid-like component. For interphase cells, we see a trend of decreasing elastic moduli and cortical tension in dependence of increasing malignant potential (Fig. 1c and Fig. S1). By contrast, for mitotic cells, we find the opposite trend of increasing elastic moduli and cortical tension in dependence of increasing malignant potential (Fig. 1e and Fig. S1). Corresponding loss tangents *G*″/*G*′ for the cortex of interphase and mitotic cells are presented in panels Fig. 1d,f. Solid lines describe model loss tangents as inferred by Eqn. (2) and (3) using parameters from fits to measured storage and loss moduli as shown in Fig. 1c,e. Unlike previously reported [23], we do not see an increase of the loss tangent of cells in dependence of malignant potential, see Fig. 1d,f. Boxplots of cortical stiffness quantified by |*G*\*| and phase shift tan^−1^(*G*″/*G*′) measured at a frequency of 1 Hz as well as active cortical tension are presented in Fig. S1a-c.

Frequency-dependence of storage and loss moduli are well described by a rheological model that anticipates a constant relaxation spectrum up to a slowest cut-off relaxation time scale (see Eqn. (2) and (3) in Materials and Methods). Model fits to averaged storage and loss moduli are depicted as solid lines in (Fig. 1c,e). Corresponding fitting parameters *τ_max_*, *K_h_* and *η* are shown in the caption of Fig. 1. Model parameters obtained from fitting of individual rheological curves are shown in Table I and Fig. S1e-g (*τ_max_* quantifies the slowest relaxation time scale, *K_h_* rates the mechanical strength of relaxation modes and *η* captures a purely viscous contribution that captures an increase of the loss modulus at high frequencies). Decreasing values of elastic moduli with malignant potential are reflected by a corresponding decrease of the parameter *K_h_* in interphase. By contrast, an increase of *K_h_* with malignant potential is observed in mitosis. Variations of the parameters *τ_max_* and *η* point to additional differences in cytoskeletal regulation amongst the three different cell lines such as different turnover time of cortical constituents and variation in cytoplasmic viscosity. In conclusion, we observe here a trend of decreasing interphase cortical moduli and increasing mitotic cortical moduli in conjunction with increasing malignant potential in three breast epithelial cell lines.

**Table I.** Rheological parameters obtained from fits of individual cells of two breast-epithelial cell lines with epithelial phenotype: MCF-10A and MCF-7, and one with mesenchymal-like phenotype: MDA-MB231, (data set as in Fig. 1c-f). Values represent medians of elastic modulus *K_h_*, slowest relaxation time *τ_max_* and viscous contribution *η* as determined by fitting of Eqn. (2) and (3) (see Materials and Methods). Furthermore, we report median values of active cortical tension *γ_act_* and acquired normalised confinement cell height 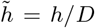 during the measurement. Here, *D* is the estimated cell diameter of the unperturbed cell. P-values refer to comparison with data obtained for mesenchymal-like cell line MDA-MB231.

| Cell type | $K_h$ (mN/m) | p-value | $\tau_{max}$ (s) | p-value | $\eta$ (Pa s) | p-value | $\gamma_{act}$ (mN/m) | p-value | $\tilde{h}$ | n(#) |
| --- | --- | --- | --- | --- | --- | --- | --- | --- | --- | --- |
| MCF-10A interphase | 5.11 | 0.0014 | 9.66 | 0.039 | 0.31 | 0.003 | 0.51 | 0.003 | $0.64 \pm 0.02$ | 32 |
| MCF-7 interphase | 4.43 | 0.020 | 8.89 | 0.1 | 0.41 | 0.0001 | 0.41 | 0.041 | $0.64 \pm 0.02$ | 39 |
| MDA-MB231 interphase | 3.10 | 1 | 10.25 | 1 | 0.01 | 1 | 0.33 | 1 | $0.64 \pm 0.02$ | 24 |
| MCF-10A mitosis | 6.13 | 0.0001 | 23.24 | 0.056 | 0.43 | 0.004 | 0.4 | 0.006 | $0.61 \pm 0.02$ | 32 |
| MCF-7 mitosis | 11.23 | 0.0001 | 9.45 | 0.26 | 0.32 | 0.0001 | 0.78 | 0.36 | $0.61 \pm 0.02$ | 38 |
| MDA-MB231 mitosis | 18.65 | 1 | 11.69 | 1 | 0.93 | 1 | 0.61 | 1 | $0.61 \pm 0.02$ | 41 |

EMT is known to be a hallmark of cancer progression and malignancy [6, 31]. Therefore, we wanted to test to which extent actin cortex rheology is influenced by EMT-associated cortex remodelling. Our previous study reported EMT-induced cell-cycle dependent changes of cortical mechanics in breast epithelial cells sampled at a frequency of 1 Hz [10]. However, a complete characterisation of cortical rheological changes upon EMT is lacking. To this end, we mechanically sampled the actin cortex of MCF-7 and MCF-10A cells before and after pharmacologically induced EMT (Fig. 2). EMT was induced by TPA- or TGF-*β*1-treatment as described before, see [10] and Materials and Methods and Fig. S2c. Measurements were performed on non-adherent interphase cells and cells in mitotic arrest for frequencies between 0.02 −2 Hz. Obtained estimates of cortical complex elastic moduli in dependence of frequency are shown in Fig. 2. We find that moduli of non-adherent interphase MCF-7 and MCF-10A cells are lowered upon EMT at all frequencies measured. Vice versa, complex elastic moduli increased at all frequencies in mitotic cells of both cell lines. Data points were fitted with our rheological model as given by Eqn. (2) and (3) (solid lines in all panels of Fig. 2). Corresponding fitting parameters *τ_max_*, *K_h_* and *η* are shown in the caption of Fig. 2. The loss tangents showed no EMT-related trend in mitotic and interphase MCF-10A and MCF-7 cells (Fig. S2a-b). In addition, as reported previously [10], active cortical tension decreases in interphase upon EMT but shows the opposite trend in mitosis (Table II).

**Table II.**
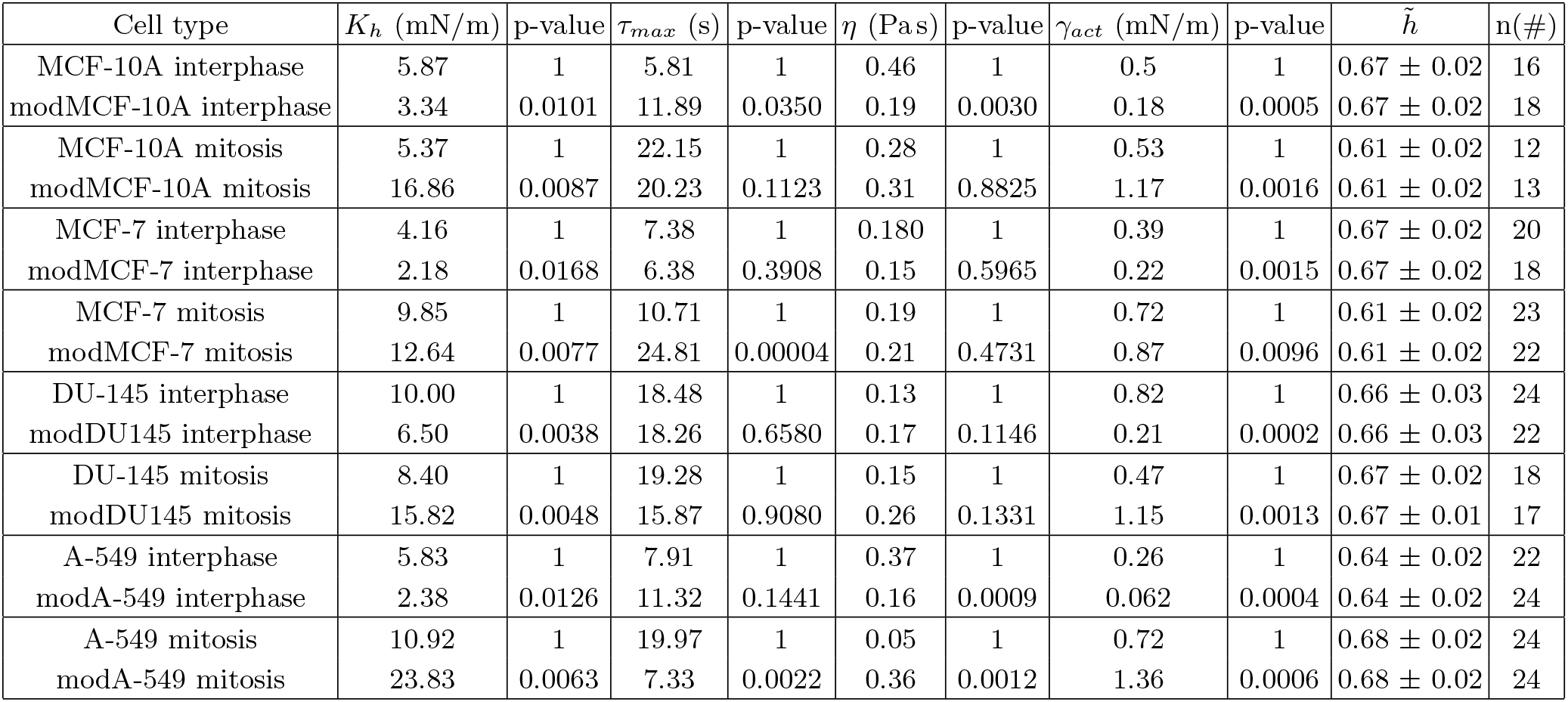
Rheological parameters obtained from fits of individual cells of different epithelial cell lines in pre- and post-EMT conditions (data sets as in Fig. 2 and 3a-d). Cells were measured in interphase in suspension or in mitotic arrest as indicated. Values represent medians of elastic modulus *K_h_*, slowest relaxation time *τ_max_* and viscous contribution *η* as determined by fitting of Eqn. (2) and (3) (see Materials and Methods). Furthermore, we report median values of active cortical tension *γ_act_* and normalised confinement cell height 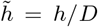 during the measurement. Here, *D* is the estimated cell diameter of the unperturbed cell.

So far, we only tested the influence of EMT on cortical mechanics in breast epithelial cells. To investigate if EMT-associated rheological changes are conserved in cells of other tissue origin, we applied our assay to the cell lines DU-145 and A-549 originating from explanted prostate carcinoma [32] and lung tumor tissue [33], respectively. Both cell lines show an epithelial phenotype in standard media. Using an established protocol, we induced EMT pharmacologically through TGF-*β*1 treatment [34, 35], see Materials and Methods. In order to verify EMT in our treated cells, we performed immunoblotting and observed the characteristic decrease in E-cadherin and a corresponding increase in N-cadherin and Vimentin (Fig. S3a-b). Furthermore, we generated immunofluorescence micrographs of E-cadherin and Vimentin before and after EMT confirming the characteristic EMT-induced changes (Fig. S3f).

As before, we measured actin cortex rheology before and after EMT in non-adherent interphase and mitotic cells obtaining complex elastic moduli as depicted in Fig. 3a-d. Analogous to breast epithelial cell lines, we find that complex elastic moduli of non-adherent interphase cells are lowered upon EMT (Fig. 3a-b) while moduli of mitotic cortices increased at all frequencies (Fig. 3c-d). Data points were fitted with our rheological model as given by Eqn. (2) and (3) (solid lines in all panels of Fig. 3). Again, we find on average good agreement with our measurements. Fitting parameters *τ_max_*, *K_h_* and *η* of averaged rheological data are shown in the caption of Fig. 3. Corresponding loss tangents showed no clear EMT-induced trend in interphase and mitotic DU-145 and A-549 cells (Fig. S3c-d). Furthermore, we present boxplots of cortical stiffness quantified by |*G*\*| and phase shift tan^−1^(*G*″/*G*′) measured at frequency of 1 Hz. Measured active cortical tensions are presented in Fig. 3e-g. Similar to breast epithelial cell lines, we find a significant EMT-induced cell mechanical trend of decreased cortical stiffness and active cortical tension in interphase DU-145 and A-549 cells and the opposite trend in mitosis (Fig. 3e-g).

For EMT-related rheological measurements on MCF-10A, MCF-7, DU-145 and A-549 (see Fig. 2, 3), rheological fitting parameters as well as active cortical tension *γ_act_* obtained for individual cells are summarised in Table II. Furthermore, Fig. 4 shows corresponding boxplots of obtained parameter distributions of rheological fitting parameters *K_h_*, *τ_max_* and *η*.

In interphase, for all cell lines considered, the modulus *K_h_* and active cortical tension decreases upon pharmacological EMT induction (Fig. 4a and Fig. 5, top). By contrast, for all cell lines, *K_h_* and active cortical tension *γ_act_* increase upon EMT in mitosis (Fig. 4a and Fig. 5, bottom).

**Figure 5.**
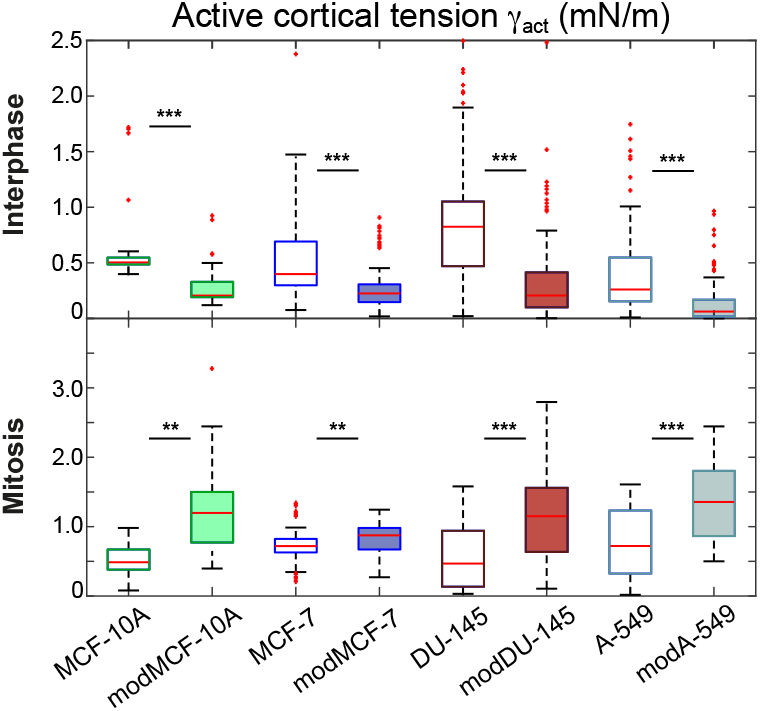
Boxplot indicating the distribution of active cortical tension *γ_act_* for measured MCF-10A, MCF-7,DU-145, A-549 interphase and mitotic cells before and after EMT (same data set as in Fig. 4, Post-EMT cells are referred to as modMCF-10A, modMCF-7, modDU-145 and mod-A-549, respectively. n.s.: *p* > 0.05, *: *p* < 0.05, **: *p* < 0.01, * * *: *p* < 0.001.)

The characteristic time scale *τ_max_* shows overall no consistent EMT-related trend (Fig. 4b); However, in mitotic MCF-7 cells, *τ_max_* increases upon EMT reflecting an EMT-triggered cortical solidification, i.e. a drop of the loss tangent *G”*/*G′* at all frequencies. We conjecture that this change in mechanical relaxation time is related to previously reported EMT-induced changes of actin cortical turnover (see Fig. S3 in [10]). Vice versa, in mitotic A-549 cells, *τ_max_* decreases upon EMT reflecting an EMT-triggered cortical fluidization in mitosis, i.e. a rise of the loss tangent *G*″/*G*′ at all frequencies. Interestingly, we find that EMT-induced changes of *τ_max_* tend to be opposite in interphase and mitosis if present. However, the direction and magnitude of the change varies between cell lines.

Also, the viscous contribution *η* shows no consistent trend upon EMT with no significant changes of measured viscosity values in most cell lines (Fig. 4c). We do, however, observe an EMT-induced viscosity decrease in MCF-10A and A-549 interphase cells and a viscosity increase in mitotic A-549 cells (Fig. 4c). These changes of *η* might point to a corresponding increase or decrease of cytoplasmic viscosity upon EMT in respective cell lines.

## Discussion

In this study, we investigated the influence of EMT on frequency-dependent actin cortex rheology of human cells in interphase and mitosis. To this end, we utilised an established AFM-based cell-confinement assay which was previously shown to provide a characterisation of the complex elastic modulus and the active tension of the actin cortex [10, 21, 22, 36]. The presence of active stresses in a material can confound the values of the elastic modulus obtained from mechanical measurements [37–39]. Our assay avoids this effect by separating active and passive cortical stresses in the data analysis, thus, reporting active cortical tension and cortical viscoelastic moduli as independent parameters [22].

Inducing EMT in epithelial cell lines derived from breast, lung and prostate tissue, we observed a general trend of a reduction of actin cortex elastic moduli in interphase cells and a decrease of active cortical tension. By contrast, we found that moduli of the actin cortex of mitotic cells are generally higher than those of interphase cells with a distinct increase upon EMT. Furthermore, mitotic cells develop a higher active cortical tension upon EMT, see Fig. 2 and 3. By comparison, we find that an increase in malignant potential in breast epithelial cells gives the same qualitative trend as EMT induction (reduction in interphase, increase in mitosis, see Fig. 1 and S1).

Observed cell-mechanical changes upon EMT are likely related to EMT-induced activity changes of cytoskeletal regulator proteins Rac1 and RhoA which were consistently reported by many studies [10–13]. In particular, activity changes of Rac1 can account for the cell-cycle dependent mechanical changes of the actin cortex by an as yet unknown downstream mechanism (Rac1 activity reduction causes a corresponding cell-mechanical change that is opposite in interphase and mitosis as was shown in [10]). In addition, we identified in previous work that EMT induces a myosin decrease at the interphase cortex and a myosin and actin increase at the mitotic cortex in MCF-7 cells; cortex thickness was not significantly affected by EMT [10]. Further characterisation of cortex molecular changes underlying the observed mechanical phenotype will be the object of future research.

Previously, many cell-mechanical studies have reported interphase cell softening in correlation with an increase in malignancy potential [10, 14–20, 30]. Furthermore, in mitotic cells, it was shown that Ras-activation enhances mitotic stiffness as measured by AFM indentation [30]. As EMT is a hallmark of cancer progression and malignancy, our findings are consistent with these earlier studies. Other studies have monitored cellular mechanical changes upon EMT. Osborne *et al*. observed cell-softening upon EMT in isolated adherent murine breast epithelial cells using adherent magnetic beads deflected by a magnetic tweezer [40]. Interestingly, Schneider *et al*. reported effective cell-stiffening upon EMT in adherent sheets of murine breast epithelial cells using AFM indentation at the apical cell apex [41]. Here, the authors attribute their findings to an increased actin cytoskeletal contribution in cellular force response after EMT rather than to stiffening of the actin cytoskeleton itself. A corresponding reorganisation and redistribution of actin cytoskeleton upon EMT was shown by Haynes *et al*. [42]. Our previous study on breast epithelial cells also showed a softening of interphase and a stiffening of mitotic cells’ cortices upon EMT measured at a single frequency of 1 Hz. Furthermore, it was shown that EMT enhances traction forces in adherent MCF-10A cells hinting at increased actomyosin contractility in stress fibres post-EMT [43]. This contractility trend in adherent interphase cells is interestingly opposite to our finding of EMT-induced contractility reduction in the actin cortex of non-adherent interphase cells. This discrepancy underscores the differences in actin cytoskeletal regulation in the two different actin structures. In fact, inhibition of the actin nucleator Arp2/3 was shown to reduce traction forces in adherent cells [44] while Arp2/3 inhibition was shown to increase contractility in the actin cortex [45, 46]. As EMT has been associated with a strong activity increase of Rac1 [10–13], a major activator of Arp2/3, we speculate that the opposite contractility response to Arp2/3 activity is at the heart of different EMT-induced contractility phenotypes in the actin cortex and in actin stress fibers.

In summary, we showed in this study that measured actin cortex mechanics is captured by a universal rheological model described by a constant relaxation spectrum of time scales up to a slowest relaxation time scale *τ_max_* which we determined to be in the range of 5 – 25 s and which is on the same order of magnitude as the turnover of cortical cytoskeletal components [3]. Our data suggest that this rheological model is conserved for cells from different tissue origin and throughout the cell-cycle. The cancer-related cellular transformation EMT leads to a roughly universal downscaling of viscoelastic moduli of the cortex in interphase and an opposite upscaling of moduli in mitosis as reflected by corresponding changes of the model parameter *K_h_*. Concomitantly, active cortical tensions decrease in interphase and increase in mitosis upon EMT. We conclude that these EMT-induced rheological changes likely reflect major changes in the organisation, structure and functionality of the actin cortex upon EMT.

## Materials and Methods

### Cell culture

The cultured cells were maintained as follows: MCF-7, DU-145 and A-549 cells were grown in RPMI-1640 medium (PN:2187-034, life technologies) supplemented with 10% (v/v) fetal bovine serum, 100 *μ*g/mL penicillin, 100 μg/ml streptomycin (all Invitrogen) at 37°C with 5% CO_2_ MCF-10A cells were cultured in DMEM/F12 medium (PN:11330-032, Invitrogen) supplemented with 5% (v/v) horse serum (PN:16050-122, Invitrogen), 100 *μ*g/mL penicillin, 100 *μ*g/mL streptomycin (all Invitrogen), 20 mg/mL epidermal growth factor (PN:AF-100-15, Peprotech), 0.5 mg/mL hydrocortisone (PN:H-0888, Sigma), 100 ng/mL cholera toxin (PN:C-8052, Sigma), 10 *μ*g/mL insulin (PN:I-1882, Sigma) at 37°C with 5% CO_2_. In MCF-7 cells, EMT was induced by incubating cells in medium supplemented with 100 nM 12-O-tetradecanoylphorbol-13-acetate (TPA) (PN:P8139, Sigma) for 48 hours prior to measurement [47]. In MCF-10A, EMT was induced by incubating cells in medium supplemented with 10 ng/mL TGF-*β*1 (PN:90900-1, BPSBioscience, San Diego, USA) one week prior to measurement [48–50]. For DU-145 and A-549, EMT was induced by 24 hours treatment with 5 ng/mL TGF-*β*1 [34, 35]. EMT changed the cellular phenotype in culture notably. In control conditions, epithelial cells grow in dense islands of several cells while after induced EMT cells grow more isolated from each other.

### Experimental setup

To prepare mitotic cells for AFM measurements, ≈10 000 cells were seeded in a cuboidal silicon cultivation chamber (0.56 cm^2^ area, from cutting ibidi 12-well chamber) that was placed in a 35 mm cell culture dish (fluorodish FD35-100, glass bottom, World Precision Instruments) one day before the measurement so that a confluency of ≈30% was reached on the measurement day. Mitotic arrest by supplementing S-trityl-L-cysteine (STC, Sigma) was induced 2 - 8 h before the measurement at a concentration of 2 *μ*M. For measurement, mitotically arrested cells were identified by their shape. Their uncompressed diameter ranged typically from 18 to 23 *μ*m.

To prepare AFM measurements of suspended interphase cells, cell culture dishes (fluorodish FD35-100) were coated by incubating the dish at 37 °C with 0.5 mg/mL Poly(L-Lysine)-PEG dissolved in PBS for 1 h (PLL(20)-g[3.5]-PEG(2), SuSoS, Dubendorf, Switzerland) to prevent cell adhesion. Prior to measurements, cultured cells were detached by the addition of 0.05% trypsin-EDTA (Invitrogen). Approximately 30 000 cells in suspension were placed in the coated culture dish.

The culture medium was changed to CO2-independent DMEM (PN: 12800-017, Invitrogen) with 4 mM NaHCO_3_ buffered with 20 *μ*M HEPES/NaOH pH 7.2, for AFM experiments ≈2 h before the measurement [10, 21, 22, 51].

The experimental setup included an AFM (Nanowizard I, JPK Instruments) that was mounted on a Zeiss Axiovert 200M optical, wide-field microscope using a 20x objective (Zeiss, Plan Apochromat, NA = 0.8) along with a CCD camera (DMK 23U445 from Theimagingsource). Cell culture dishes were kept in a Petri dish heater (JPK Instruments) at 37 °C during the experiment. Prior to every experiment, the spring constant of the cantilever was calibrated by thermal noise analysis (built-in software, JPK) using a correction factor of 0.817 for rectangular cantilevers [52]. The cantilevers used were tipless, 200 - 350 *μ*m long, 35 *μ*m wide, and 2 *μ*m thick (NSC12/tipless/no aluminium or CSC37/tipless/no aluminium, Mikromasch). The nominal force constants of the cantilevers ranged between 0.2 and 0.8 N/m. The cantilevers were supplied with a wedge, consisting of UV curing adhesive (Norland 63, Norland Products Inc.) to correct for the 10° tilt [53]. The measured force, piezo height, and time were output at a time resolution of 500 Hz.

### Dynamic AFM-based cell confinement

Preceding every cell compression, the AFM cantilever was lowered to the dish bottom in the vicinity of the cell until it touched the surface and then retracted to approximately 14 *μ*m above the surface. Subsequently, the free cantilever was moved and placed on top of the cell. Thereupon, a bright-field image of the equatorial plane of the confined cell is recorded in order to evaluate the equatorial radius *R_eq_* at a defined cell height *h*. Cells were confined between dish bottom and cantilever wedge. Then, oscillatory height modulations of the AFM cantilever were carried out with oscillation amplitudes of 0.25 *μ*m in a frequency range of 0.02 and 2 Hz. We verified that rheological measurements are in the linear range for this amplitude choice, see Fig. S5 in the Supporting Material and Fig. S3 in [22]

During this procedure, the cell was on average kept at a normalised height 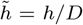 between 60 – 70%, where *D* = 2(3/(4*π*)*V*)^1/3^ and *V* is the estimated cell volume. Using molecular perturbation with cytoskeletal drugs, we could show in previous work that at these confinement levels, the resulting mechanical response of the cell measured in this setup is dominated by the actin cortex, see Fig. 4 in [22] and Fig. S7 in [10]. This is further corroborated by our observation that the smallest diameter of the ellipsoidal cell nucleus in suspended interphase cells of all cell lines under consideration is smaller than 60% of the cell diameter, see Fig. S4. Additionally, it has been shown that for a nucleus-based force response, when measuring cells in suspension with AFM, a confinement of more than 50% of the cell diameter is needed [54].

It is noteworthy that the measured cortical elastic modulus increases with cell confinement height (up to 30% in the confinement height range of 60 – 70%). This phenomenon can be attributed to varying contributions of area shear and area dilation during cortical deformation as was pointed out by our previous study [36]. Therefore, for each experiment, different conditions were measured on the same day with the same average cell confinement height (±3%) and the same AFM cantilever. Average normalised height values adopted for each experiment are shown in Table I and II.

### Estimating cellular shape and volume during AFM measurements

For our analysis, we anticipate that confined cells adopt droplet shapes of minimal surface area as reported in [21]. Cell volume was estimated by imaging the equatorial plane of a confined cell at retracted AFM cantilever, i.e. of a cell height h of ≈ 14 *μ*m. The cell volume was determined by the approximative formula 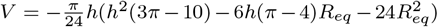 [22, 36]. For measurement analysis, the cell volume was anticipated to be constant during the measurement of the cell [21, 22]. Further, we estimated radii of principle curvatures *R*_1_ and *R*_2_ at the equator of the cell surface as *h*/2 and *R_eq_* [22]. The area of contact between cells and cantilever was estimated as 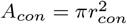 with the contact radius *r*_con_ determined by the approximative formula 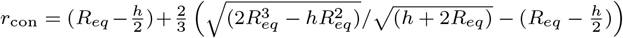 as described in [21].

### Data analysis

In our analysis, the force response of the cell is translated into an effective cortical tension 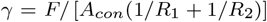, where *A_con_* is the contact area between confined cell and AFM cantilever and *R*_1_ and *R*_2_ are the radii of principal curvatures of the free surface of the confined cell [10, 21, 22]. Oscillatory force and cantilever height readouts were analysed in the following way; for every time point, effective cortical tension *γ* and surface area strain (*A*(*t*) – 〈*A*〉)/〈*A*〉 were estimated. An amplitude and a phase angle associated to the oscillatory time variation of effective tension *γ* and surface area strain are extracted by sinusoidal fits. To estimate the value of the complex elastic modulus at a distinct frequency, we determine the phase angles *φ_γ_* and *φ_ϵ_*A*__* as well as amplitudes *A_γ_* and *A_ϵ_*A*__* of active cortical tension and surface area strain, respectively. The complex elastic modulus at this frequency is then calculated as 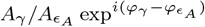.

Model Equations (2) and (3) were fitted to data of complex elastic moduli with a least square fit in Matlab using the command lsqcurvefit. Statistical analyses of cortex mechanical parameters were performed in Matlab using the commands ‘boxplot’ and ‘ranksum’ to generate boxplots and determine p-values from a Mann-Whitney U-test (two-tailed), respectively.

### Model of cortical force response

In steady state, at constant cantilever height, the obtained tension value *γ* is approximately constant in time and interpreted as the active cortical tension *γ_act_*. If the cortical film is subject to dynamic deformation, the mechanical response of the cortex changes over time. We assume a linear viscoelastic response of the form [22, 55]

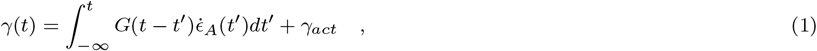

where *ϵ_*A*_*(*t*) = (*A*(*t*) – *A*_0_)/*A*_0_ is the area strain, *A*_0_ is a reference area, and the dot indicates a time derivative. The relaxation modulus *G*(*t*) shows the time-dependent response, which consists of memory effects due to viscoelastic material properties.

Measurement results obtained from mitotic HeLa cells put forward a relaxation spectrum *h*(*τ*) = *K_h_* for *τ* ≤ *τ_max_* and *h*(*τ*) = 0 for *τ* > *τ_max_* for cortical material [22]. The corresponding relaxation modulus reads *G*(*t*) = –*K_h_*,*Ei*(–*t*/*τ*_max_) where *Ei*(*z*) is the exponential integral function [22]. The associated complex elastic modulus *G*\*(*ω*) is calculated as 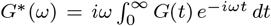. Adding a viscous contribution *η* to capture the viscous response of the cytoplasm or the medium in particular at high frequencies, we obtain the following functional form of storage and loss moduli

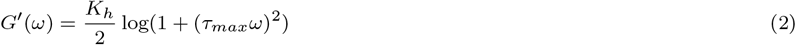

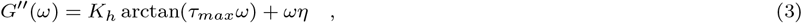

where *G*\* = *G*′ + *iG*′. Eqn. (2) and (3) were used to perform the frequency-dependent fits in this paper. They constitute a simple model to discuss the rheology of the cortical layer based on a characteristic time scale τ_max_ and a complex elastic modulus *K_h_*.

We find that all our rheological measurements of the actin cortex are reasonably well captured by this simple rheological model, see solid lines in Fig. 1, 2 and 3. This model is consistent with the idea that the turnover of F-actin and different actin cross-linkers contributes multiple cortical relaxation time scales (namely a continuous spectrum up to a longest relaxation time scale *τ_max_*) [56, 57]. Naturally, cortical relaxation time scales must be bounded by the time scale of turnover of F-actin in the cortex. In fact, our estimated range for the largest relaxation time scale *τ_max_* ≈ 5-25s agrees reasonably well with actin turnover time scales predicted by photobleaching experiments at the cortex [3, 10, 58].

### Western blotting

Protein expression in DU-145 and A-549 before and after EMT was analysed using Western blotting. Cells were seeded onto a 6-well plate and grown up to a confluency of 80-90% with or without EMT-inducing agents. Thereafter, cells were lysed in SDS sample/lysis buffer (62.5 mM TrisHcl pH 6.8, 2% SDS, 10% Glycerol, 50 mM DTT and 0.01%Bromophenolblue). Cell lysates were incubated for 30 mins with the lysis buffer at 4°C. Subsequently, they were sonicated for 2 mins to break-down undissolved components and then boiled for 10 minutes. 10 - 20 μL of the cleared lysate was then used for immunoblotting. The cleared lysates were first run on precast protein gels (PN:456-1096 or 456-1093, Bio-Rad) in MOPS SDS running buffer (B0001, Invitrogen). Subsequently, proteins were transferred to nitrocellulose membranes (Merck). Nitrocellulose membranes were blocked with 5% (w/v) skimmed milk powder (T145.1, Carl Roth, Karlsruhe, Germany) in TBST (20 mM/L Tris-HCl, 137 mM/L NaCl, 0.1% Tween 20 (pH 7.6)) for 1 h at room temperature followed by washing with TBST, and incubation at 4°C overnight with the corresponding primary antibody diluted 1:100 (Vimentin), 1:1000 (E-Cadherin), 1:500 (N-Cadherin) or 1:3000 (GAPDH) in 1% (w/v) bovine serum albumin/TBST solution. Thereupon, the blots were incubated with appropriate secondary antibodies conjugated to horseradish peroxidase, Goat anti-mouse HRP (PN: ab97023, Abcam) or Goat anti-rabbit HRP (PN: ab97051, Abcam) at 1:3000 - 1:5000 dilution in 5% (w/v) skimmed milk powder in TBST for 1 h at room temperature. After TBST washings, specifically bound antibodies were detected using Pierce enhanced chemiluminescence substrate (ECL) (PN:32109, Invitrogen). The bands were visualised and analysed using a CCD-based digital blot scanner, ImageQuant LAS4000 (GE Healthcare Europe, Freiburg, Germany). Primary antibodies used are as follows: E-Cadherin (PN:60335-1-lg, Proteintech), N-Cadherin (PN:13116T, Cell Signaling Technology), Vimentin (PN:AMF-17b, Developmental Studies Hybridoma Bank, Iowa City, IA), GAPDH (PN:ab9485, Abcam).

### Immunofluorescence

The adherent cells were fixed with 3.7% formaldehyde/PBS for 15 min at RT, followed by a 15 min permeabilisation step in 0.2% Triton X-100 at RT. The samples were then blocked with 3% BSA/PBS for 1h at RT. The cells were then stained overnight at 4°C with 1:40 Vimentin (PN:AMF-17b, Developmental Studies Hybridoma Bank, Iowa City, IA) or 1:100 E-Cadherin (PN:60335-1-lg, Proteintech) primary antibodies. The samples were then treated with 1:1000 secondary Alexa Fluor 488 (PN:A32723, Invitrogen) and 5 *μ*g/mL DAPI in 3% BSA/PBS solution for 2 h at RT. Images were recorded using a Zeiss LSM700 confocal microscope of the CMCB light microscopy facility, incorporating a Zeiss C-Apochromat 40x NA 1.2 water objective.

### Nuclei imaging

For nuclei size imaging, suspended interphase cells were prepared as for AFM measurements. Approximately 30 mins before imaging, 0.1 *μ*g/mL of Hoechst 3342 (PN:62249, Invitrogen) was added to the samples in order to stain the cell nucleus. Exemplary images are shown in Fig. S4.

## Acknowledgments

We thank Marta Urbanska and Jochen Guck for provision of the cell line A-549. Further, we thank Isabel Richter, Ivan Antonenko and Rewati Limaye for support of western blotting work. E.F.-F. acknowledges financial support from the Deutsche Forschungsgemeinschaft, projects FI 2260/4-1 and FI 2260/5-1. K.H. and E.F.-F. were further supported by the Deutsche Forschungsgemeinschaft under Germany’s Excellence Strategy, EXC-2068-390729961, Cluster of Excellence Physics of Life of TU Dresden.

## Author Contributions

K.H. and E.F.-F. designed the research. K.H. and A.F. performed experiments. K.H. and E.F.-F. carried out the data analysis. K.H. and E.F.-F. wrote the manuscript.

