## Supplementary Information for "EMT changes actin cortex rheology in a cell-cycle dependent manner"

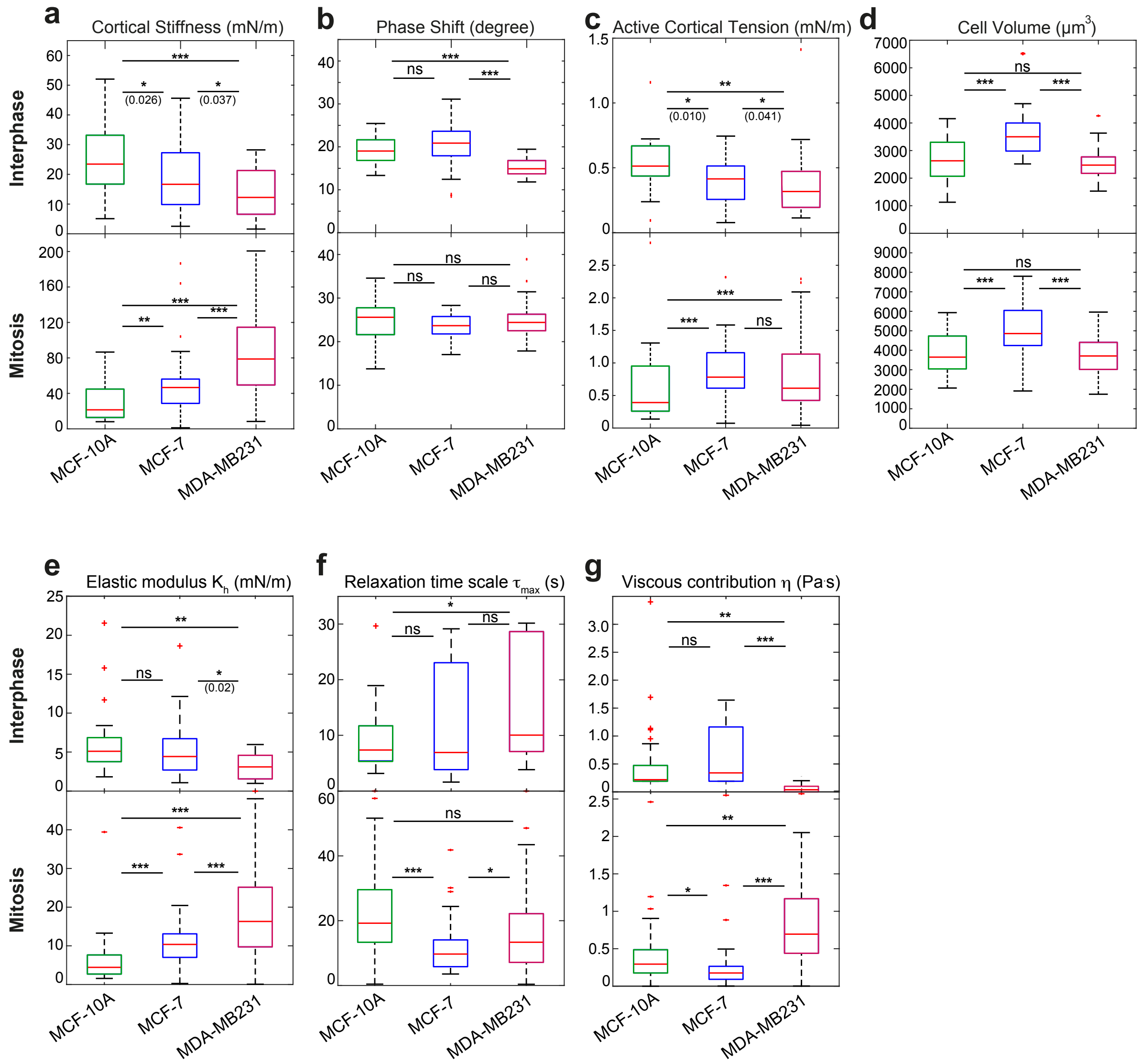


Figure S1. a-d) Cortical stiffness $|G^*|$ (a), phase shift (b), active cortical tension (c) and cell volume (d) measured for MCF-10A, MCF-7 and MDA-MB231 suspended interphase cells (top row) and cells in STC-arrested mitosis (bottom row) at 1 Hz frequency corresponding to measurements in Fig. 1, main text. e-g) Boxplots indicating the distribution of rheological fitting parameters (e) $K_h$, (f) $\tau_{max}$ and (g) $\eta$ for measured MCF-10A, MCF-7 and MDA-MB23 interphase and mitotic cells corresponding to measurements in Fig. 1, main text. Number of cells analysed (a-e): interphase: MCF-10A n=32, MCF-7 n=39, MDA-MB231 n=24, mitosis: MCF-10A n=32, MCF-7 n=38 and MDA-MB231 n=41. n.s.: $p > 0.05, * : p < 0.05, ** : p < 0.01, *** : p < 0.001$. Normalised confinement heights chosen during respective measurements are given in Table 1, main text.

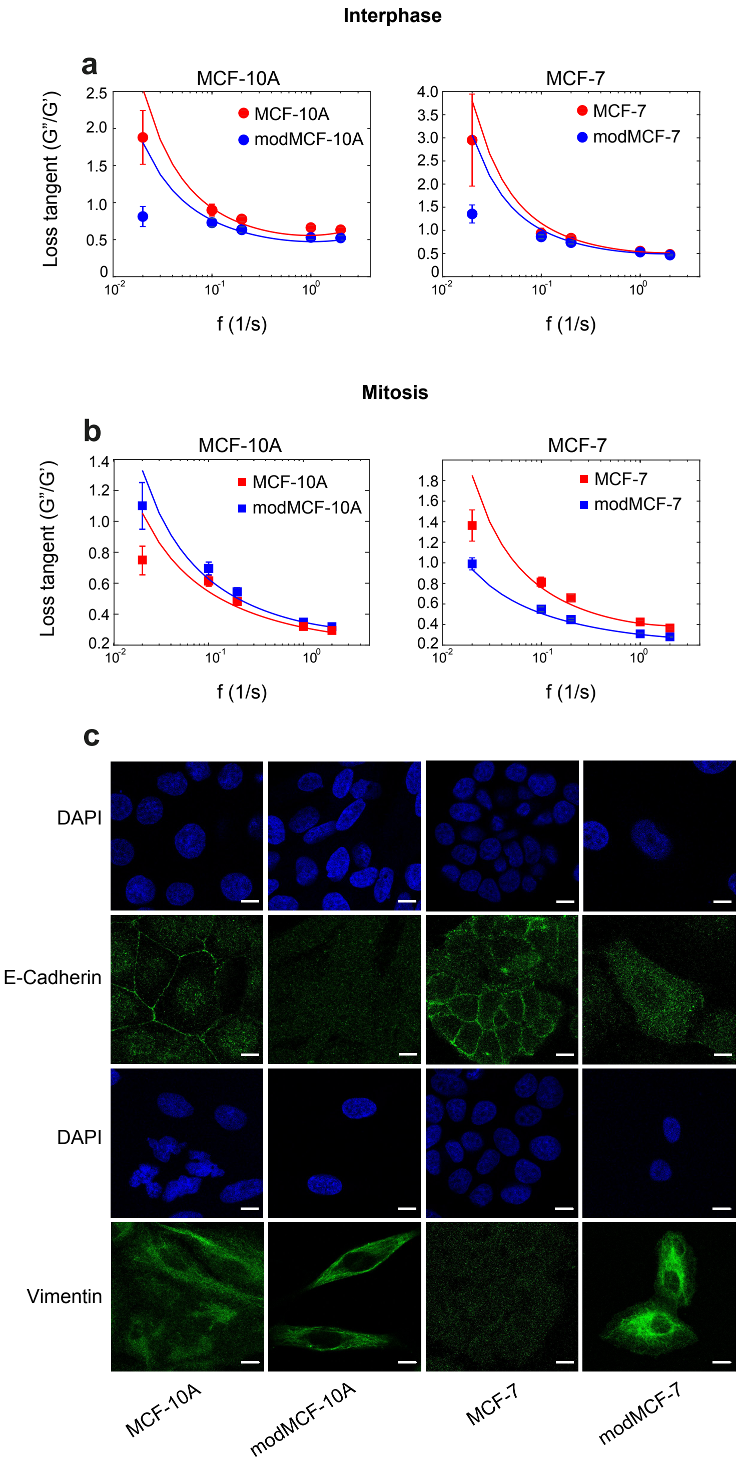


Figure S2. a-b) Loss tangents of MCF-10A and MCF-7 before and after EMT in suspended interphase (a) and STC-arrested mitotic cells (b) corresponding to measurements in Fig. 2, main text. Post-EMT cells are referred to as modMCF-10A and modMCF-7, respectively. Number of cells analysed: interphase: MCF-10A n=16, modMCF-10A n=18, MCF-7 n=20, modMCF-7 n=18, mitosis: MCF-10A n=21, modMCF-10A n=23, MCF-7 n=23 and modMCF-7 n=22. Error bars indicate standard error of the mean. c) Immunofluorescence of E-Cadherin and Vimentin in adherent cells before and after EMT in MCF-10A and MCF-7. Scale bar: 10 $\mu$m.

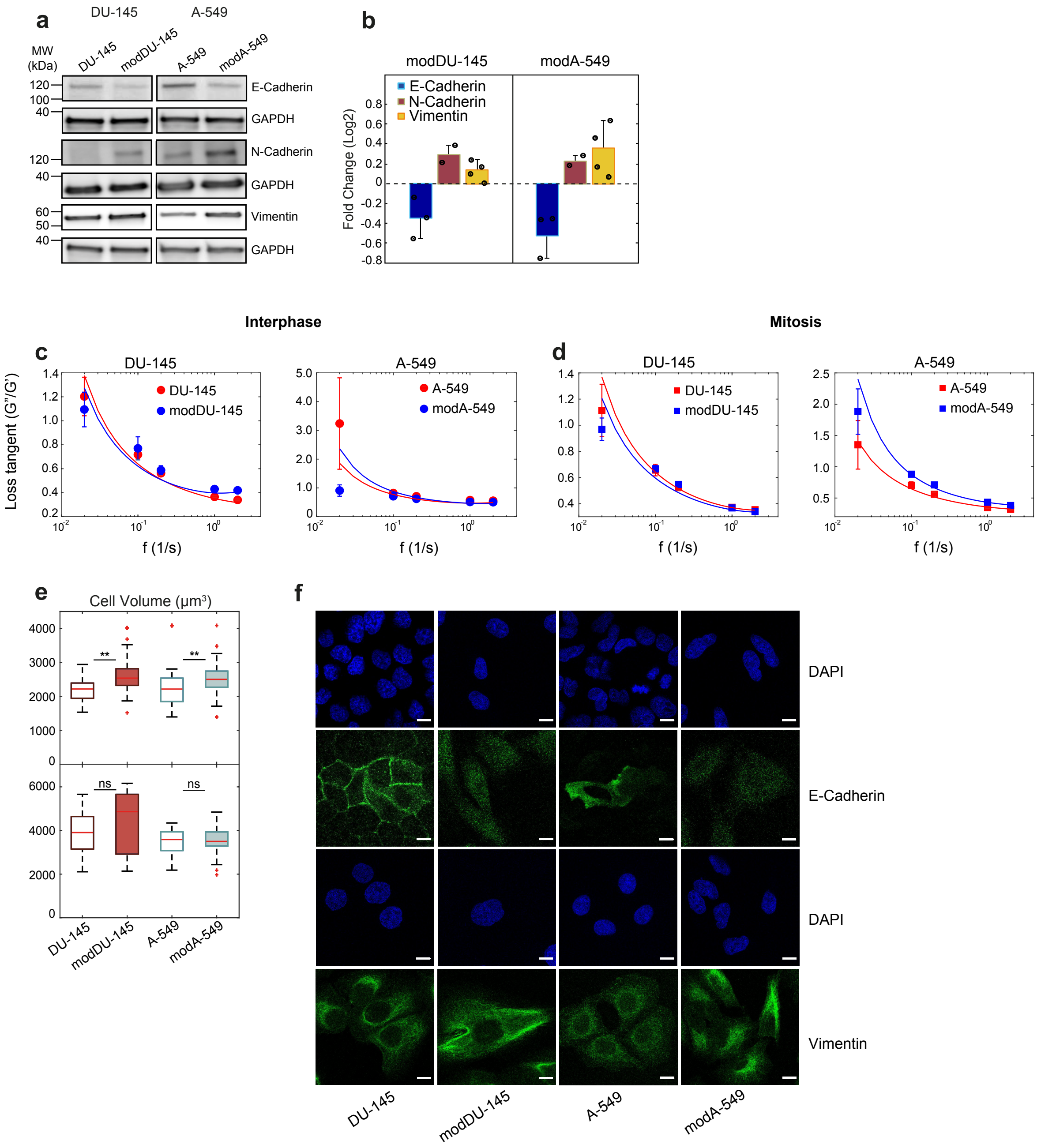


Figure S3. a-b) Protein expression changes upon TGF-$\beta$1-induced EMT (5 ng/mL for 24 hours treatment) in DU-145 and A-549 cells: a) Western blots showing E-Cadherin (top row), Vimentin (third row) and N-Cadherin (fifth row) before and after TGF-$\beta$1-induced EMT. b) Quantification of relative changes of E-Cadherin, N-cadherin and Vimentin from Western blots. Quantifications were normalised by GAPDH bands. Error bars represent standard deviations. Individual data points are shown with black dots. c-d) Loss tangents of DU-145 and A-549 before and after EMT in suspended interphase (c) and cells in mitotic arrest (b) corresponding to measurements in Fig. 3, main text. Error bars indicate standard errors of the mean. e) Cell volumes of suspended interphase (top row) and STC-arrested mitotic (bottom row) DU-145 and A-549 cells before and after EMT corresponding to measurements in Fig. 3, main text. Post-EMT cells are referred to as modDU-145 and modA-549, respectively. Number of cells analysed (a-b): E-Cadherin n=3, N-cadherin n=2 and Vimentin n=4. (c-d): interphase: DU145 n=24, modDU145 n=22, A-549 n=22, modA-549 n=24, mitosis: DU145 n=18, modDU145 n=17, A-549 n=24 and modA-549 n=24, (e): interphase: DU145 n=36, modDU145 n=36, A-549 n=35, modA-549 n=39, mitosis: DU145 n=34, modDU145 n=34, A-549 n=34 and modA-549 n=29. n.s.: $p > 0.05, * : p < 0.05, ** : p < 0.01, *** : p < 0.001$. f) Immunofluorescence of E-Cadherin and Vimentin in adherent cells before and after EMT in DU-145 and A-549. Scale bar: 10 $\mu$m.

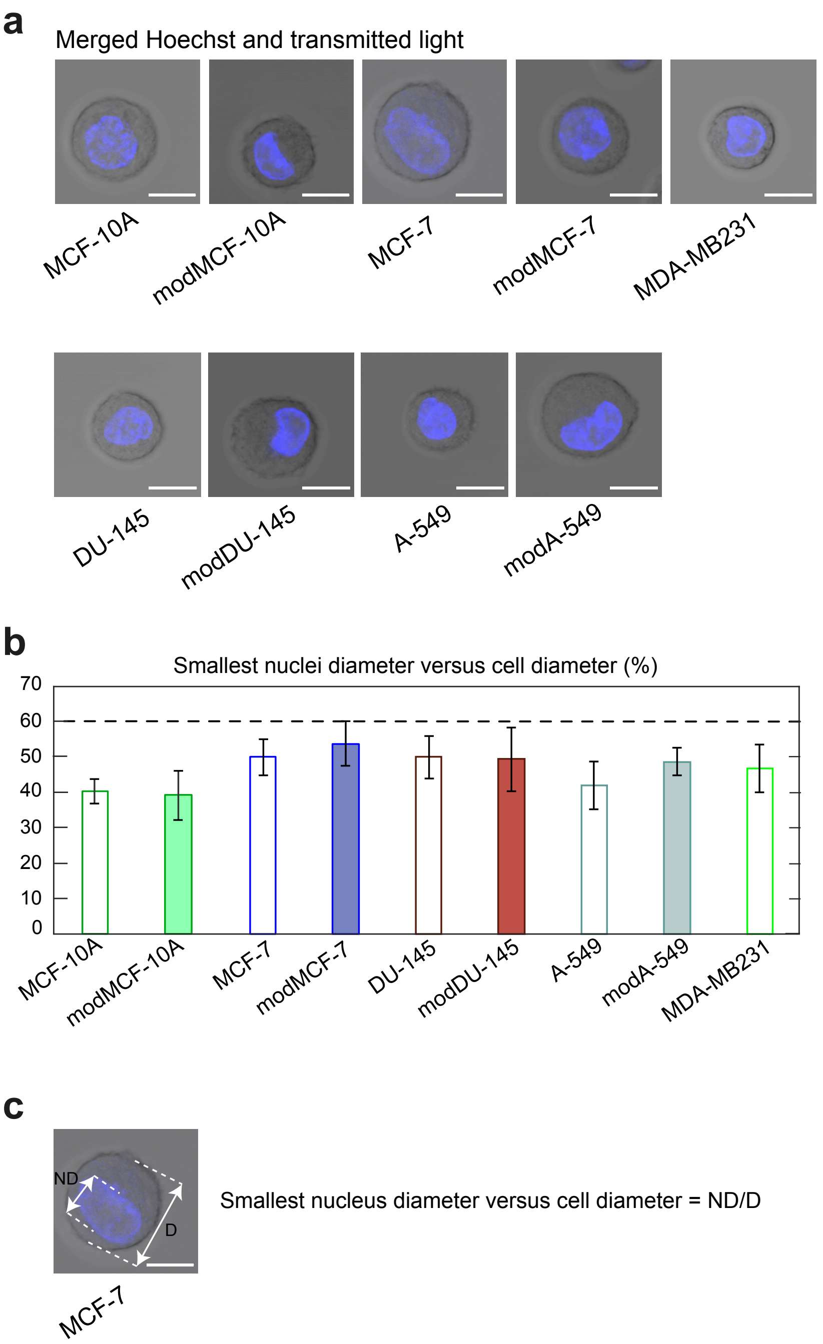


Figure S4. Extension of cell nuclei in different cell lines. a) Exemplary pictures of suspended interphase cells of different cell lines under consideration before and after EMT. The nucleus is shown in blue (Hoechst staining). Scale bar: 10 $\mu$m. b) Quantification of smallest nucleus diameter relative to the cell diameter (quantification as indicated in panel c). c) Exemplary quantification of smallest nucleus diameter relative to the cell diameter. Scale bar: 10 $\mu$m. Post-EMT cells are referred to as modMCF-10A and modMCF-7, modDU-145 and modA-549, respectively. Number of cells quantified: MCF-10A n=20, modMCF-10A n=20, MCF-7 n=24, modMCF-7 n=22, MDA-MB231 n=20, DU-145 n=20, modDU-145 n=20, A-549 n=20 and modA-549 n=20. Error bars indicate standard deviations.

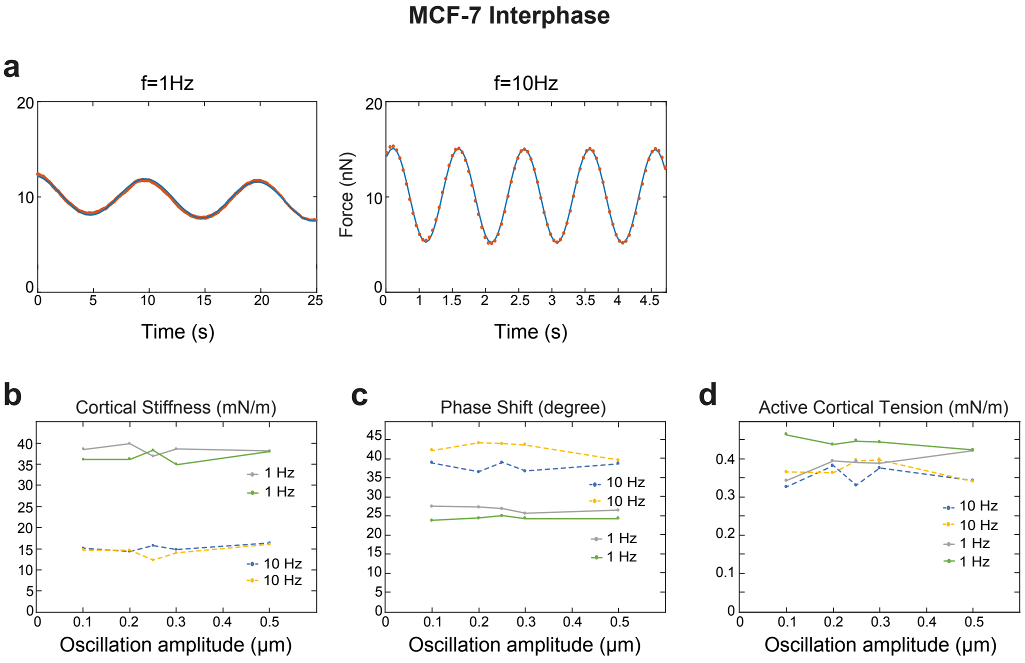


Figure S5. Force oscillations are sinusoidal and scale linear with oscillation amplitudes. a)Exemplary force oscillations at our standard cantilever height oscillation amplitude of $0.25\,\mu$m at frequencies $f = 1\,$Hz and $f = 10\,$Hz. Force oscillations are close to sinusoidal in shape in the measured frequency range (data points shown in orange dots, sinusoidal fit in blue line). b-c) Measured cortical stiffnesses, phase shifts and cortical tensions in dependence of cantilever height oscillation amplitude imposed during dynamic cell confinement. Obtained mechanical parameters are largely independent of the imposed oscillation amplitude up to height amplitude of 0.5 $\mu$m indicating that we operate in the regime of linear viscoelasticity.
